# Symbiont-Screener: a reference-free filter to automatically separate host sequences and contaminants for long reads or co-barcoded reads by unsupervised clustering

**DOI:** 10.1101/2020.10.26.354621

**Authors:** Mengyang Xu, Lidong Guo, Chengcheng Shi, Xiaochuan Liu, Jianwei Chen, Xin Liu, Guangyi Fan

## Abstract

Decontamination is necessary for eliminating the effect of foreign genomes on the symbiont studies and biomedical discoveries. However, direct extraction of host sequencing reads with no references remains challenging. Here, we present a trio-based method to classify the host error-prone long reads or sparse co-barcoded reads prior to assembly, free of any alignments against DNA or protein references. This method first identifies high-confident host reads by haplotype-specific *k*-mers inherited from parents, and then groups remaining host reads by the unsupervised clustering. Experimental results demonstrated that this approach successfully classified up to 97.38% of the host human long reads with the precision rate of 99.9999%, and 79.95% host co-barcoded reads with the precision rate of 98.36% using an artificially mixed data. Moreover, the tool also exhibited a good performance on the decontamination of the real algae data. The purified reads reconstructed two haplotypes and improved the assembly with larger contig NGA50 value and less misassemblies. Symbiont-Screener can be freely downloaded at https://github.com/BGI-Qingdao/Symbiont-Screener.

## Introduction

With the development and optimization of biotechnology and big data analytics, genome sequences become significant biological basis and genetic resources of modern life science. The accuracy of assembled genome sequences determines the quality of downstream applications, including gene regulation, epidemiological investigation, genome-wide association studies, and species evolutionary^1^. However, a certain number of genome assemblies, even those which have been submitted to public databases may contain foreign sequences from other species due to the impurity in raw reads^2,3^. The decontamination in the final assembly of target species is required to filter out erroneous genetic properties and eliminate their effects on further research conclusions.

On the other hand, the investigation of symbiotic or parasitic relationships between host and microbiome also requires the separation and reconstruction of genomes of all the associated species and strains. Projects such as Human Microbiome Project (HMP)^4^ and Global Catalogue of Microorganisms Project (GCM)^5^ aim to decipher the roles of microbes in host health and wellbeing through metagenomic analysis of associated microbiota. The studies of metagenomics also require the separation of host and foreign reads with high accuracy and completeness.

The sources of DNA contamination in the genomes are complex. One possibility refers to the mixture of organisms in the biological sample. Most organisms in nature, in fact, live with symbionts, parasites and commensals, and it is technically difficult to separate metazoan hosts and their attached species. Those foreign genomes can be normally found in all individuals of the same species under similar environment. Laboratory pollution or artificial experimental errors may also lead to the contamination, which possibly pollute a few of individuals by some chance.

Recently, there have been several experimental and bioinformatic approaches to remove contaminants without consideration of their sources. Embryonic DNA of target species has been reported to be extracted to avoid symbiotic contamination^6^, but it needs additional culture for embryogenesis and cannot eliminate endosymbiotic microorganisms. Current bioinformatic decontamination methods are based on the disparity in statistical or biological features for different species. DNA sequencing coverage, GC composition, and oligonucleotide frequencies can be used as indicators to partition draft assemblies^7,8^. However, these statistical methods cannot resolve inherently low read coverage of sex chromosomes and heterozygous sites of autosomes^9,10^, and wide overlap of GC content ranges for bacteria and archaea^11^ and eukaryotic plants^12^ and animals^13,14^. More common approaches filter sequences based on the biological nucleotide or protein similarity of assemblies to known genomes or public databases, and provide taxonomy information^3,15–17^. However, these methods are not suitable for *de novo* projects and do not consider the redundancy or overlap among references. Thus, they risk mis-eliminating target sequences and decrease the data efficiency. Moreover, these tools need pre-assembled contigs to enhance the rate of genomic identification, and thus cannot resolve the contamination-induced misassemblies.

The technical development of single-molecule long reads (Pacific Biosciences (PacBio) and Oxford Nanopore Technologies (ONT)) and synthetic long reads or co-barcoded reads (single tube Long Fragment Reads (stLFR), 10X Genomics,...) utilizes the long-range information to resolve complicated regions in genomes such as repeats and centromeres where traditional next-generation short reads fail^18–21^. Meanwhile, the pedigree may provide unique inherent information. Recent studies have illustrated that the trio-binning strategy with long reads or co-barcoded reads enables haplotype-aware genome assembly with high accuracy and completeness, which promotes studies of genomic diseases and evolutionary relationships^22–24^. However, there are few specific decontamination tools designed for error-prone long reads or sparse co-barcoded reads, and current local-alignment-based methods originally designed for short reads or pre-assembled contigs could be less efficient and more time-consuming.

Here, we present a novel pipeline with unsupervised clustering, Symbiont-Screener, to automatically filter contamination sequences specifically designed for singlemolecule long reads or co-barcoded reads for the first time. This read classification method prior to assembly utilizes the hereditary stability of the host species, eliminates contamination according to their sources, identifies host raw reads prior to assembly in a fast and effective way, and independently *de novo* assembles the host genomes without disruption of symbiotic or parasitic contamination. The *in silico* method classifies raw reads and resolves the contamination-induced assembly errors at the source. The raw reads are marked by the host’s characteristic *k*-mers rather than alignments, making it more suitable for *de novo* projects without references. More importantly, we hierarchically apply the trio information and specifically remove targeted contaminants according to their sources by generating characteristic markers. For a single long read or barcoded long fragment with sufficient long-range information, those markers can judge it based on the sparse variant sites and avoid the problem of genomic similarity among different species. Using Symbiont-Screener to separate human chromosome 19 (Chr19) and eight individual bacteria species, we obtained host long reads with a precision rate of 99.9999% and a recall rate of 97.38%, and a precision of 98.36% and a recall of 79.95% for the co-barcoded reads, respectively.

## Results

### Construction of genome sequences

We applied our Symbiont-Screener on an empirical genome dataset consisting of the host chromosome and the contamination of a defined microbial mock community. The host denotes to one of the vertebrate human chromosomes, Chr19, isolated from a male sample. We categorized all the microbial species into four groups according to their living circumstances: OC: exist with the offspring only; POC: exist with the offspring and his father; MOC: exist with the offspring and his mother; and SC: exist with all host individuals. The last group, SC, is supposed as a stable symbiont, parasite and commensal, while the other three are as accidental pollutions due to the incomplete sterilization during the library construction, for instance. Two bacterial species are assigned to each group with various sequencing depth to simulate the real situation. We marked each long reads or co-barcoded reads for each genome for downstream evaluation.

### Algorithm and designs

The design of Symbiont-Screener focuses on the stepwise purification of host sequences based on the source of foreign genomes and grouping them with high efficiency. Theoretically, the haplotype-specific markers inherited from his parents enable the identification and reconstruction of the host chromosomes, and meanwhile SC contaminants can be filtered out since their *k*-mers will no longer occur in the parental-specific marker libraries as a result of set difference operation. The difference set, however, cannot remove parental-specific contaminants, POC and MOC. On the other hand, the intersect operation of reads from all individuals ascertains the host autosomes alongside with shared contaminants SC. Nevertheless, the foreign species in POC or MOC will be discarded by the set intersection. At last, OC reads cannot be marked by none of *k*-mers, and thus being eliminated. In principle, the combination of *k*-mer set operations for three mixed samples has the ability to accurately remove all the contaminants by Algorithm 1. However, the recall rate of host reads can be limited by the heterozygosity of host and sequencing errors of input data.

**Algorithm 1.**
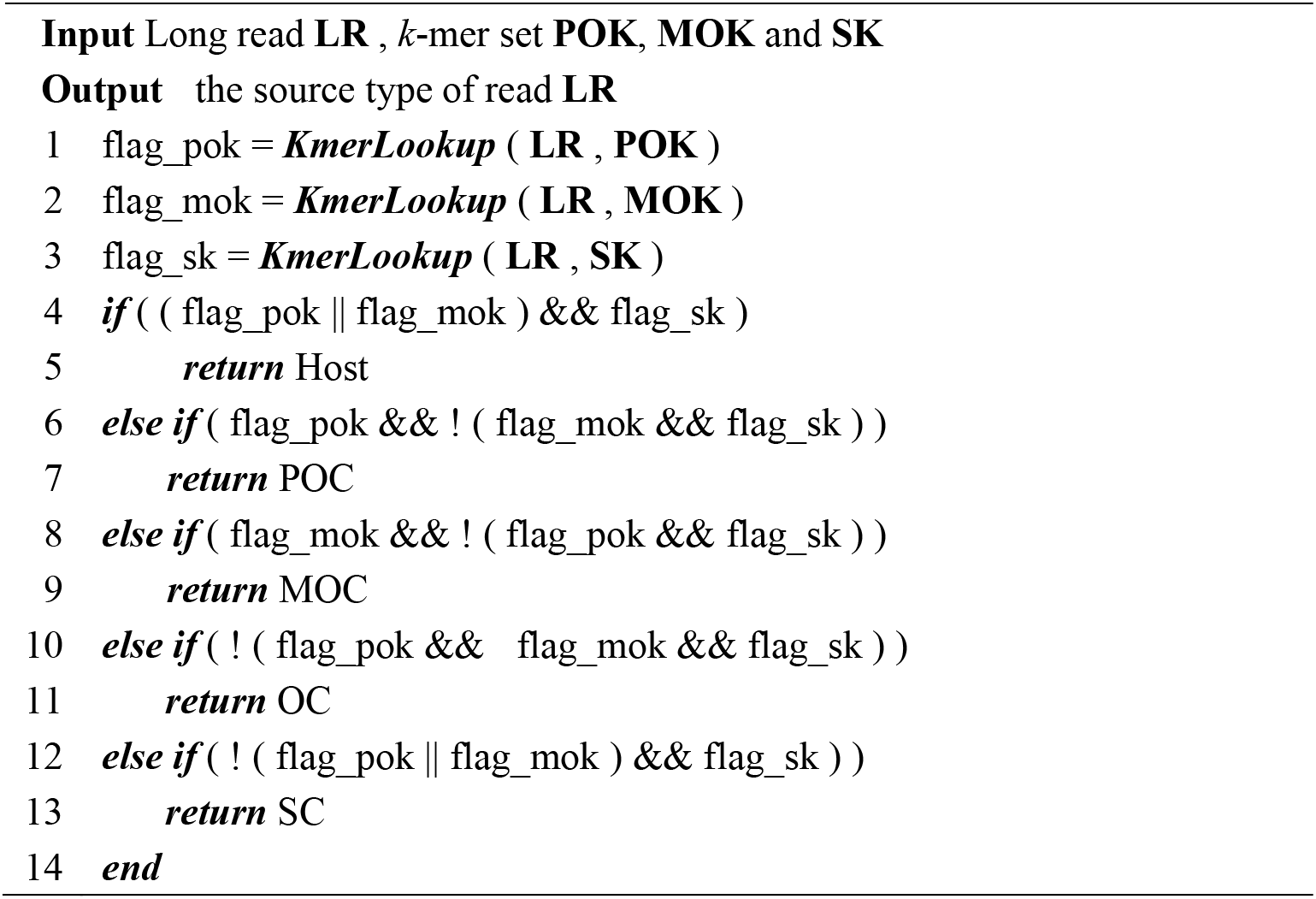
ReadSourceType

Supervised and unsupervised classification methods for pre- or post-assembled contigs/ scaffolds have been developed for metagenomic binning but can be used for purifying eukaryotic sequences^25–28^. In this study, we applied unsupervised gaussian mixture model (GMM) to cluster more reads for the host to bypass the requirement of reference sequences. In addition to the characteristic markers, commonly used statistics information can also distinguish species as complementary. In practice, the *k*-mer set operations, as well as the species differentiation in GC content and oligonucleotide frequencies consist of the main features of the algorithm (Fig. 1). Thus, the procedure of decontamination is equivalent to the read clustering based on the Mahalanobis distances to centers in the 20-dimensional feature space.

**Fig. 1.**
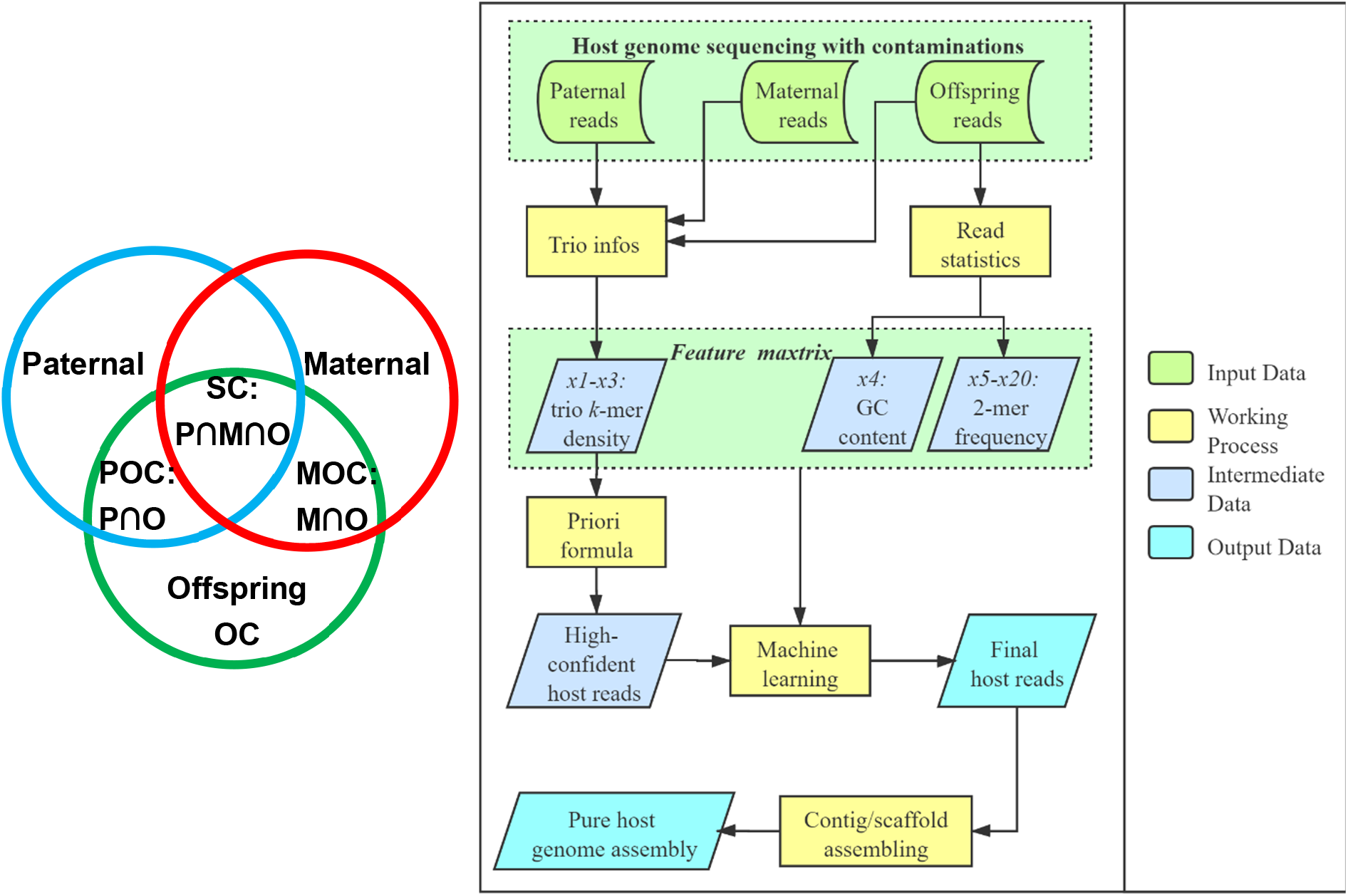
A schematic of Symbiont-Screener workflow. The left Venn diagram illustrates the categorization of contaminations and simulated datasets.

One of the problems for this unsupervised approach of Bayesian Gaussian Mixture model (BGMM) is that the clustering results were unable to identify which read group belonged to the host. However, we could use the priori high-confident host reads selected by Algorithm 1 to mark the correct group. Another problem is the instability of recall rate for multiple runs as the performance of BGMM depends on the random hits of local optimal solutions. To address this issue, we individually ran the clustering multiple times, and then merged the clustering results.

### Decontamination of long reads

There were totally 299,120 PacBio circular consensus sequence (CCS) long reads for the host Chr19 with ~50× genome coverage in the test dataset. In addition, we generated 122,088 long reads for contaminants at different sequencing coverages. 30× coverage of whole-genome sequencing (WGS) short reads for each parent sample with its corresponding foreign genomes were used to build *k*-mer libraries. The set difference operations produced 139,062 paternal-only and 169,003 maternal-only *k*-mers, respectively; while the intersect operation produced 41,876,292 shared *k*-mers among all three. With stringent thresholds, the trio-based information identified up to 146,012 priori host long reads with a precision of 99.52% but a low recall rate of 48.84% due to the relatively low heterozygous rate of the host species as well as the sequencing errors (Fig. 2(D, E, F), in black).

**Fig. 2.**
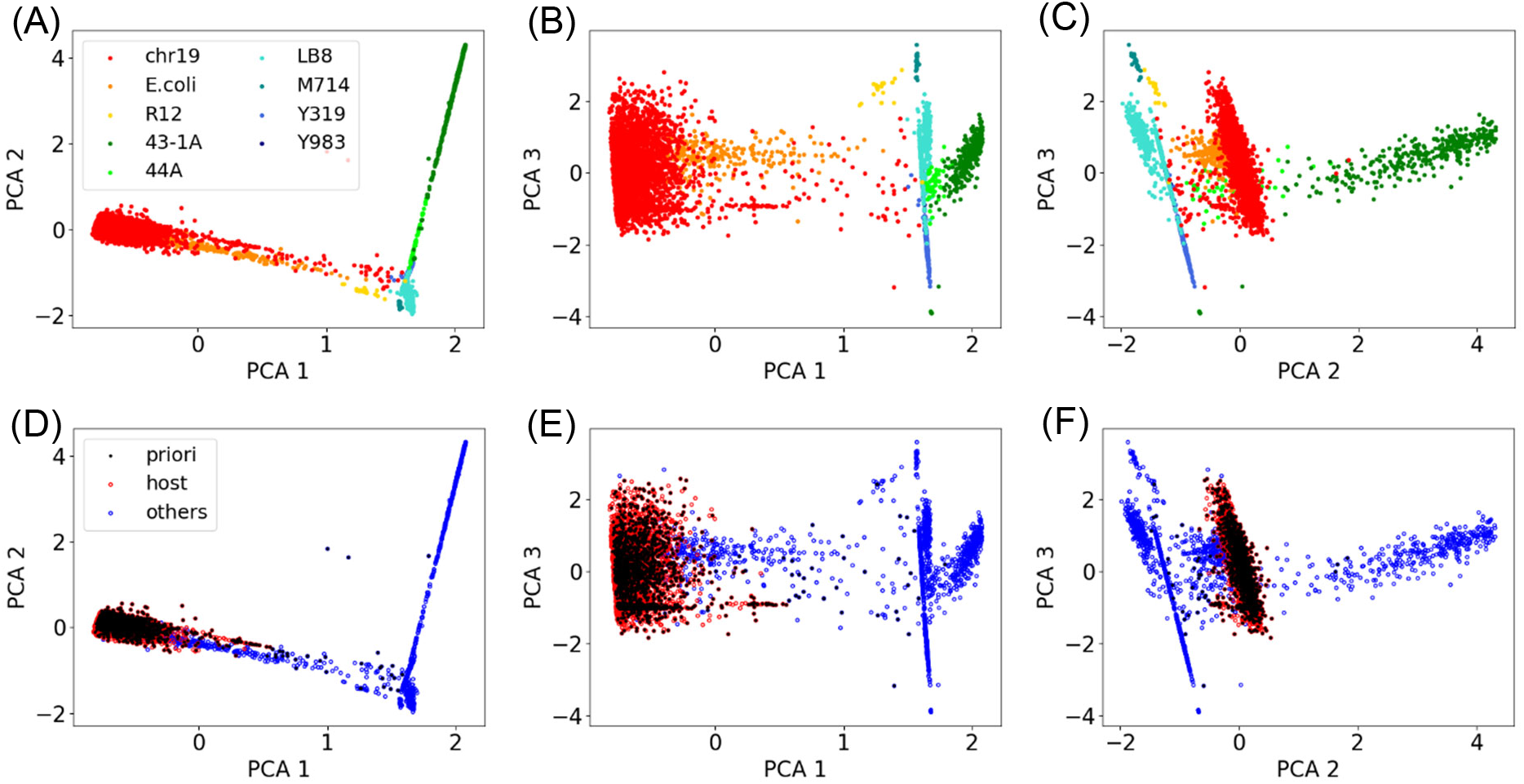
The unsupervised clustering of host reads in 20-dimensinal space. We arbitrarily selected 5000 long reads to cancel image blurring. Each point refers to a long read in the six figures. Long reads in the three top figures (A, B, C) were colored by their origins that were marked before mixing, while those in bottom (D, E, F) were colored by clustering results via Symbiont-Screener. The x, y, z axes refer to the first three most important features after principal component analysis (PCA), which contain >98% of information in total. (A, D), (B, E) and (C, F) present the clustering from different viewpoints, respectively.

We grouped more host reads by learning the features of detected priori high-confident reads. BGMM extracted 291,289 final consensus reads, of which the recall rate was greatly improved to 97.38% and the precision was stabilized at 99.9999% (Fig.2(D, E, F), in red). In addition, Fig.2 illustrated that the host group in the bottom was extended with large overlaps of the marked Chr19 reads in the top. We also note that a minority of priors was finally excluded because of their scattered distances in the feature space.

### Decontamination of co-barcoded reads

The stLFR co-barcoded reads sharing the same barcode were expected as the segments on the same long DNA molecule. There were totally 4,732,562 long fragments and correspondingly 112,788,488 co-barcoded reads in the mixed sequences, among which 78,633,158 reads (~140×) with 3,289,795 barcodes belonged to the host Chr19. The remaining long fragments were from eight bacteria with the same sequencing coverage ratios against the host as long-read dataset. The same pedigree information with set operations contributed to the priori high-confident host long fragments and constituted the first few features of machine learning. As a result, the trio-based information provided 40,626,452 priori host stLFR reads with 451,383 barcodes. The precision was 99.996% due to the more accurate base-calling in comparison with long reads. In spite of relatively low recall rate of 13.72% for long fragments, there were correspondingly 51.67% co-barcoded short reads classified as host due to the larger possibility of mapping characteristic *k*-mers for longer fragments.

In contrast to the certain number of bases in a single PacBio or ONT long reads, the number of reads tagged with the same barcode was defined as the characteristic feature, which depended on the long fragment length and read’s capture capability. All other features remained the same. The unsupervised clustering finally identified 63,912,626 final consensus co-barcoded reads with 974,269 barcodes for the host, of which the recall rate was apparently improved to 79.95% and the precision was stabilized at 98.36%.

### *k*-mer based evaluation of decontamination

To assess the effect of decontamination in the read set on the final assembly of the host, we generated the distinct 21-mers for host’s and contaminants’ references and mapped them to the clustered reads. The distinct *k*-mers were denoted as nonrepeating canonical *k*-mers in each reference assembly, and only hundreds of them were shared by different species (<0.001%). The *k*-mer completeness and depth indicate the ability to reconstruct the host genome. For the long-read experiment, the classified host long reads obtained up to 98.44% of the 42,872,525 Chr19 distinct *k*-mers regardless of repeats and non-ACGT bases (Fig. 3(A)). The high completeness along with the sufficient depth of 62.04 × ensured the high quality of final assembly. In contrast, all of eight bacteria presented a low *k*-mer completeness (<0.1%) with insufficient depth (< 0.2×) in the raw reads, which were unable to support the following genome assembly.

**Fig. 3.**
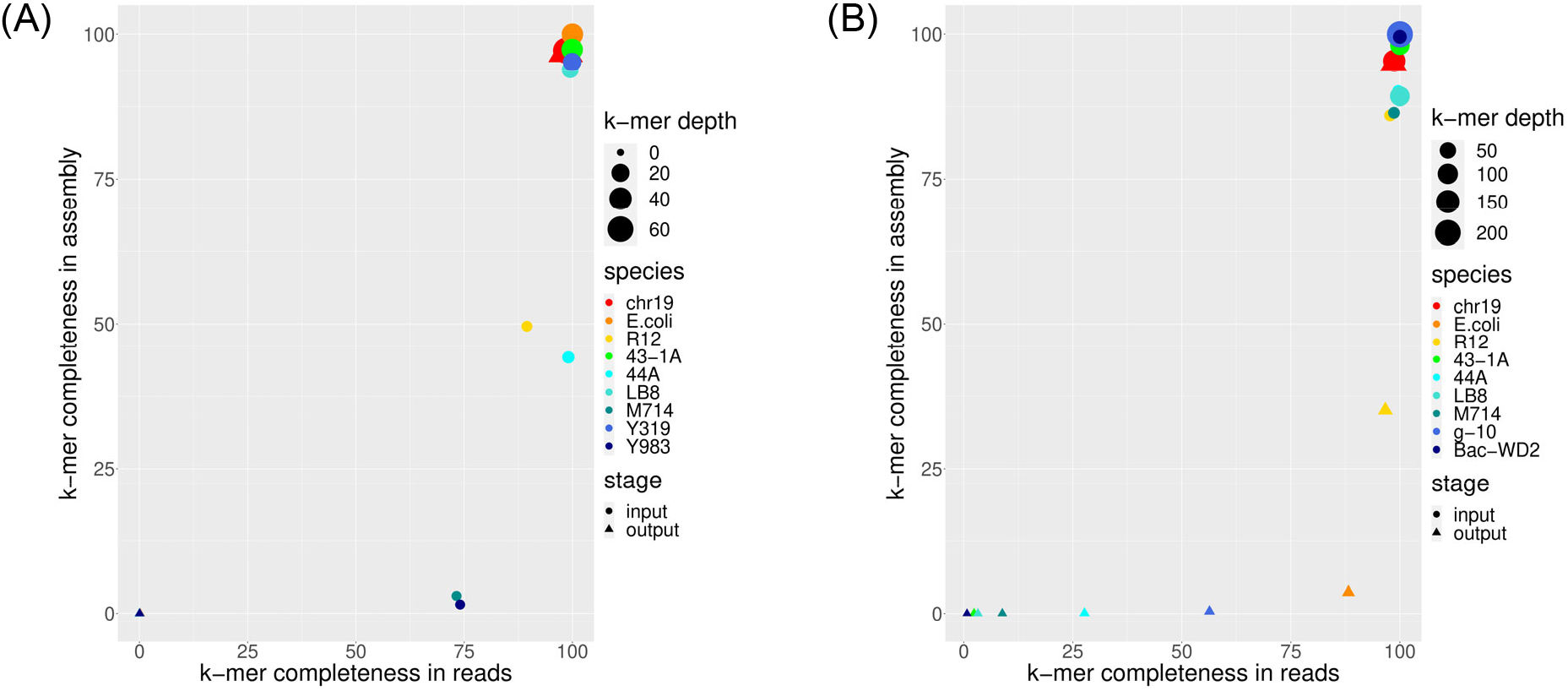
The blob plots of *k*-mer-based evaluation before and after decontamination. Host and contaminants in the (A) long-read and (B) co-barcoded-read experiments were colored by their species. Rounds and triangles denote the data of mixed sample before and after decontamination, respectively. The x axis refers to the *k*-mer completeness in raw reads while y axis refers to the *k*-mer completeness in the final assembly. The blob size is proportional to the corresponding *k*-mer depth in reads.

The co-barcoded-read experiment also showed high completeness and depth for the host reads, which successfully recovered 94.93% of the chromosome after decontamination (Fig. 3(B)). For the contaminants, only E. coli and R12 remained >80% of *k*-mers in spite of low depth (<15×). This is because the relatively greater significance of shared characteristic *k*-mers to the clustering misled the identification of their reads although they were not marked by any prior host reads. Reads for other bacteria were cleaned more thoroughly and only <1% of *k*-mers were retained in the end.

### Genome assembly of pure host

we further assembled the classified long reads and co-barcoded reads, respectively, and compared to the assemblies of mixed sample. The purified long reads with high accuracy and completeness obtained an assembly of 57,327,805 bp in size, all sequences of which were uniquely mapped to the host’s chromosome, while the total size of the mixed assembly was nearly equal to the sum of host and all contaminations. The high-quality assembly recovered 98.82% of the Chr19 reference, which agreed with the 96.94% of *k*-mer completeness in Fig. 3(B). In addition, the decontamination improved the contig NGA50 from 697 kb to 738 kb, although its contig N50 was slightly reduced, which was in concordance with 27.7% fewer misassemblies (1-10kb) and 51.9% fewer local misassemblies (<1kb). It indicated that the trio-based unsupervised clustering avoided the contamination-induced assembly errors. The single-base level accuracy was also enhanced with fewer mismatches, small insertions and deletions due to the higher error rate of mis-assembled contaminations.

The GC-coverage plots also showed obvious improvement after decontamination. The assembled contigs for the purified host eliminated suspicious peaks with apparently high or low GC content corresponding to the foreign genomes in Fig. 4(A). In addition, the decontamination extracted two clear peaks in read coverage for the diploid organism: homozygous and heterozygous in Fig. 4(B).

**Fig. 4.**
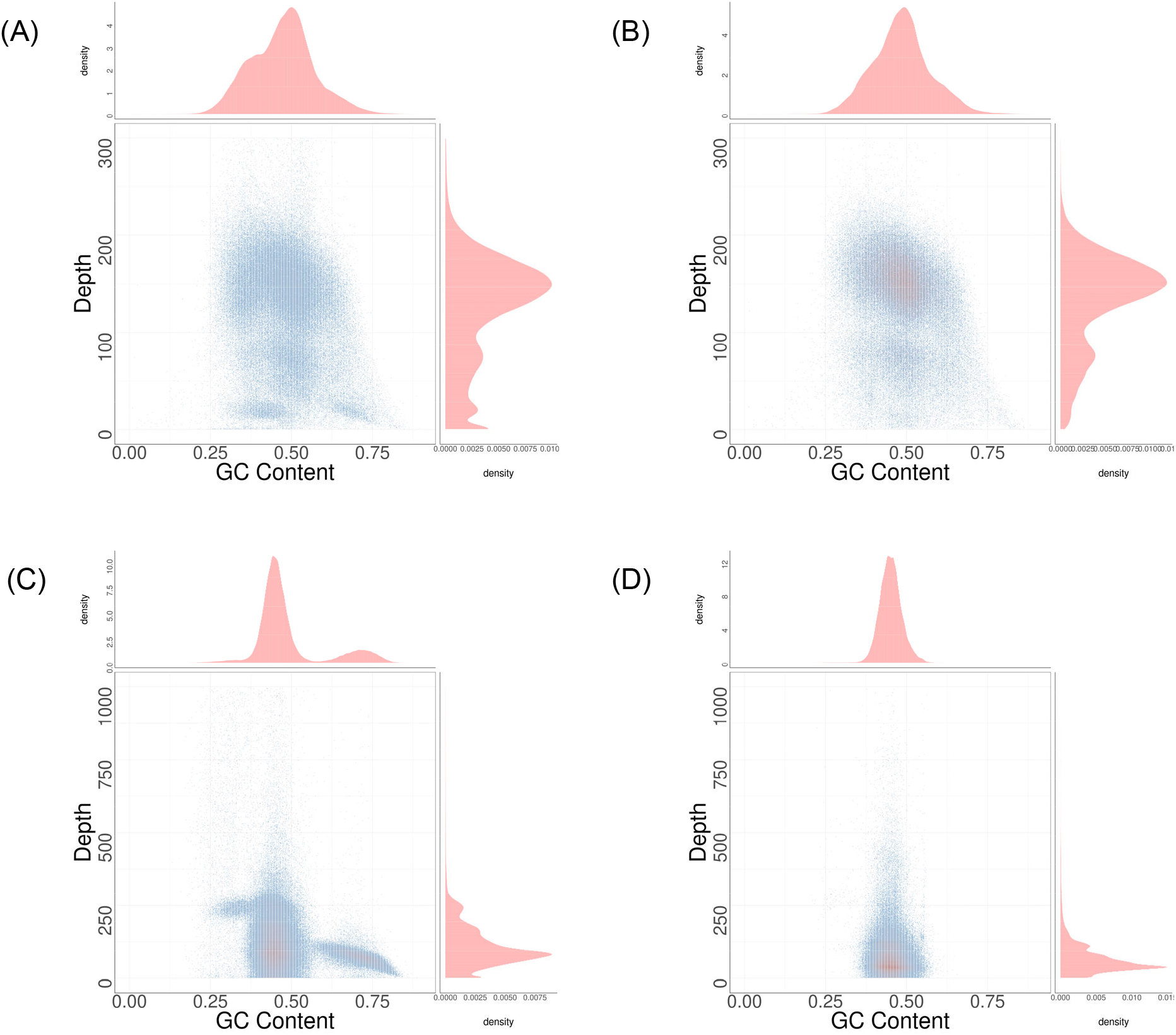
GC-depth plots for long-read assembly. Top plots refer to the assemblies of empirical long-read dataset mixed with all contaminations **(A)** and those filtered by Symbiont-Screener **(B)**, respectively. **(C)** and **(D)** refer to the assemblies of the algal real dataset before and after decontamination by Symbiont-Screener. In each plot, *x*-axis is the GC content for the assembled contigs, while *y*-axis is the coverage of short reads which are mapped to the contigs. The color scale corresponds to the densities. All assemblies were chopped into 500bp bins and their GC-content and read depth of mapped short reads were calculated and plotted by custom scripts.

The co-barcoded assembly, meanwhile, reconstructed 94.88% of the Chr19 reference. The redundant scaffolds of the assembly corresponded to the remaining contaminations, in which E. coli and R12 were only recovered by 4% and 35%. The slightly reduced genome fraction and scaffold N50 compared to the mixed assembly originated from the relatively lower sequencing coverage. However, the increased scaffold NGA50 illustrated that decontamination bypassed the misassemblies due to foreign genomes and thus avoided the misleading conclusions in downstream studies.

### Application to a real dataset

The symbiosis of algae and other species is an example of studying the mechanism of trans-kingdom coevolution^30,31^. We collected an algal sample in the wild and sequenced it using the PacBio third-generation platform. It was observed that the host algal sequences were possibly polluted by the symbiotic and other environmental prokaryotes and eukaryotes. Conventional sample purification such as sterilization and data preprocessing such as sequence search through databases were applied to remove the contamination. However, multiple peaks in the GC-coverage figure of the preliminary assembly (Fig .4(C)) indicated that these methods could not eliminate all possible foreign genomes. The clear side peak at high GC content might correspond to prokaryotic bacteria.

We applied Symbiont-Screener to the decontamination for the algal species. As it tends to clonal propagation, the trio information can be barely obtained. Instead, we selected several relative species and exploited their reference assemblies or reads to uniquely marked and grouped host reads. Other statistic features remained the same for the unsupervised clustering. The classified long reads were then assembled by Canu. As a result, the purified assembly presented only one main peak with more convergent distributions in both GC ratio and short-read coverage.

## Discussion

We introduce a novel but accurate model for decontamination that classifies reliable host raw reads from the mixed sample according to the pedigree information, that is computationally efficient without the requirement of alignment or reference sequences. The software tool based on this model, Symbiont-Screener, departs from the mapping approach in order to deal with the additional challenges arising in direct clustering error-prone long reads and sparse co-barcoded reads, especially for *de novo* projects. By taking heterozygosity and sequencing coverage into account, Symbiont-Screener is able to reconstruct the host chromosome with high contiguity and accuracy, alongside with fewer contamination-induced assembly errors. Moreover, these identified host reads via unsupervised clustering with high precision and recall rates, can be further classified into haplotype-specific groups by the parental-unique markers, which facilitate the haplotype-resolved assembly, resequencing and related studies. Additionally, our method is suitable to integrate information from multiple sequencing technique for accurate assembly of the purified host via a hybrid strategy.

The utilization of trio information that follows the Mendelian rules of inheritance, is the key for decontamination. To evaluate the significance, we *k*-merized the host’s and foreign reference assemblies, and investigated the parental-unique and shared markers (Fig. 5(A)). Owing to the low heterozygosity of human, only 0.32% and 0.39% of Chr19 *k*-mers belonged to paternal- and maternal-specific, respectively, while 97.68% were considered as shared. Those parental-unique markers (Fig. 5(B)) are able to recover the two autosomes and remove the shared bacteria (E. coli and R12), provided that the long read or the barcoded long fragment contains sufficient bases to span at least one heterozygous site. Nevertheless, the foreign sequences only existing with one parent but also found in the offspring’s contaminants (43-1A, 44A, LB8 and M714), were retained. Those sequences do not contain any shared *k*-mers and can be eliminated. Y319 and Y983 are assumed as occasional contaminants, and cannot be marked by any type of characteristic *k*-mers. In theory, all foreign genomes can be excluded by trio-based information. The sequencing error of long reads and the sparse coverage of barcoded long fragments, however, suppress the accuracy and completeness of host recognition.

**Fig. 5.**
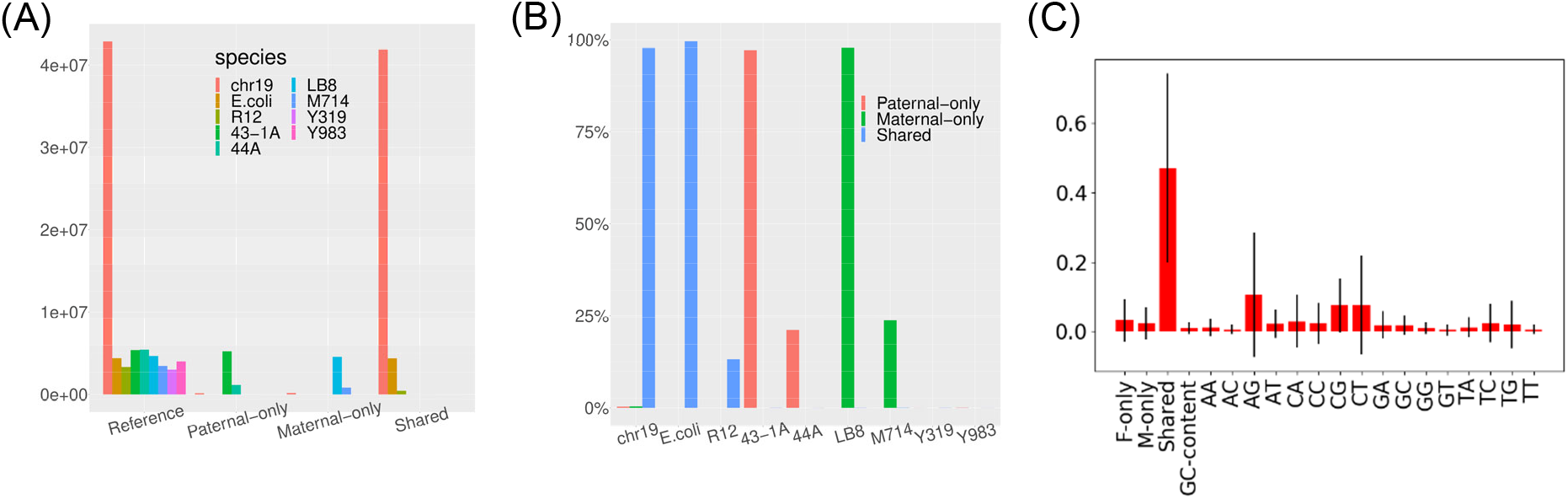
Feature importance evaluation in the clustering. (A) The *k*-mer classification of reference assemblies for the host and eight bacteria. (B) The characteristic *k*-mer component ratios of each species to illustrate the ability of hierarchical decontamination. (C) Feature importance to the clustering evaluated by random forest trees: the red bars correspond to the average significance of the feature, along with their inter-trees variability.

On the other hand, the unsupervised clustering learns features of priors including the genomic statistic information other than pedigree, and efficiently identifies more host reads thus improving the recall rate. We applied random forests of trees^32^ to evaluate the importance of features on the classification of host’s long reads. As shown in Fig. 5(C), parental-specific and shared k-mers, along with few of 2-mers are apparently more informative, and play a crucial role in the decision. Nevertheless, the requirement of trios may hamper some *de novo* projects. We are considering replacing the trio information with different individuals within the same or relative species for the decontamination.

## Methods

### Empirical genome dataset

We downloaded the PacBio CCS and stLFR sequencing data for the sample HJ from the CNGB database under accession number CNP0000091. Then all reads were aligned to the human reference hs37d5^33^ and those mapped to the chromosome 19 were extracted as the host. There were about 50× PacBio long reads and 140× stLFR co-barcoded reads left.

The assumed contaminants consist of eight single bacteria: E. coli, R12, 43-1A, 44A, LB8, M714, Y319 and Y983, among which R12, 43-1A, 44A, LB8, and M714 are unknown species isolated from soil, Y319 is identified as *Clostridium tyrobutyricum,* and Y983 belongs to *bacillus.* We sequenced ONT GidlON long reads for R12, LB8, M714, Y319 and Y983 (GridION Mk1, RRID: SCR_017986), and simulated PacBio CCS long reads for E. coli, 43-1A and 44A by PBSIM^34^ (PBSIM, RRID: SCR_002512). We also sequenced stLFR co-barcoded reads for E. coli, R12, 43-1A, 44A, LB8 and M714 using the MGIEasy stLFR Library Prep Kit on the MGISEQ-2000 platform (DNBSEQ-G400, RRID: SCR_017980). As there were no available stLFR data for Y319 and Y983, we sequenced additional two bacteria g-10 (*Serratia marcescens*) and Bac-WD2 (*Bifidobacterium longum*) as replacements. Eight species were divided into four groups of two, where E. coli and R12 belong to SC, 43-1A, and 44A belong to POC, LB8 and M714 belong to MOC, and Y319 and Y983 (g-10 and Bac-WD2) belong to OC. The former of each group has the same sequencing coverage as the host, while the latter has much lower coverages to represent the real environment (Table 1).

**Table 1.** The statistics of input reads for host and foreign species.

| Identity | Species | Genome size (bp) | # Long reads | Long-read coverage (×) | # Barcoded long fragments | # Co-barcoded reads | Long-fragment coverage (×) |
| --- | --- | --- | --- | --- | --- | --- | --- |
| Host | Chr19 | 59,128,983 | 299,120 | 50 | 3,289,795 | 78,633,158 | 140 |
| SC | E. coli | 4,502,758 | 22,417 | 50 | 2,614,807 | 5,858,854 | 140 |
|  | R12 | 3,505,947 | 1,062 | 7 | 301,546 | 636,932 | 20 |
| POC | 43-1A | 5,577,895 | 27,970 | 50 | 3,007,044 | 6,778,800 | 140 |
|  | 44A | 5,574,407 | 3,828 | 7 | 392,768 | 797,428 | 20 |
| MOC | LB8 | 4,746,099 | 37,500 | 50 | 2,480,770 | 5,630,586 | 140 |
|  | M714 | 3,481,073 | 2,150 | 7 | 313,030 | 635,792 | 20 |
| OC | Y319 (g-10.1) | 3,086,463<br>(5,216,581) | 23,326 | 50 | 3,530,874 | 13,232,512 | 140 |
|  | Y983 (Bac-WD2) | 4,059,445<br>(2,349,231) | 3,835 | 7 | 273,473 | 584,426 | 20 |

**Table 2.**
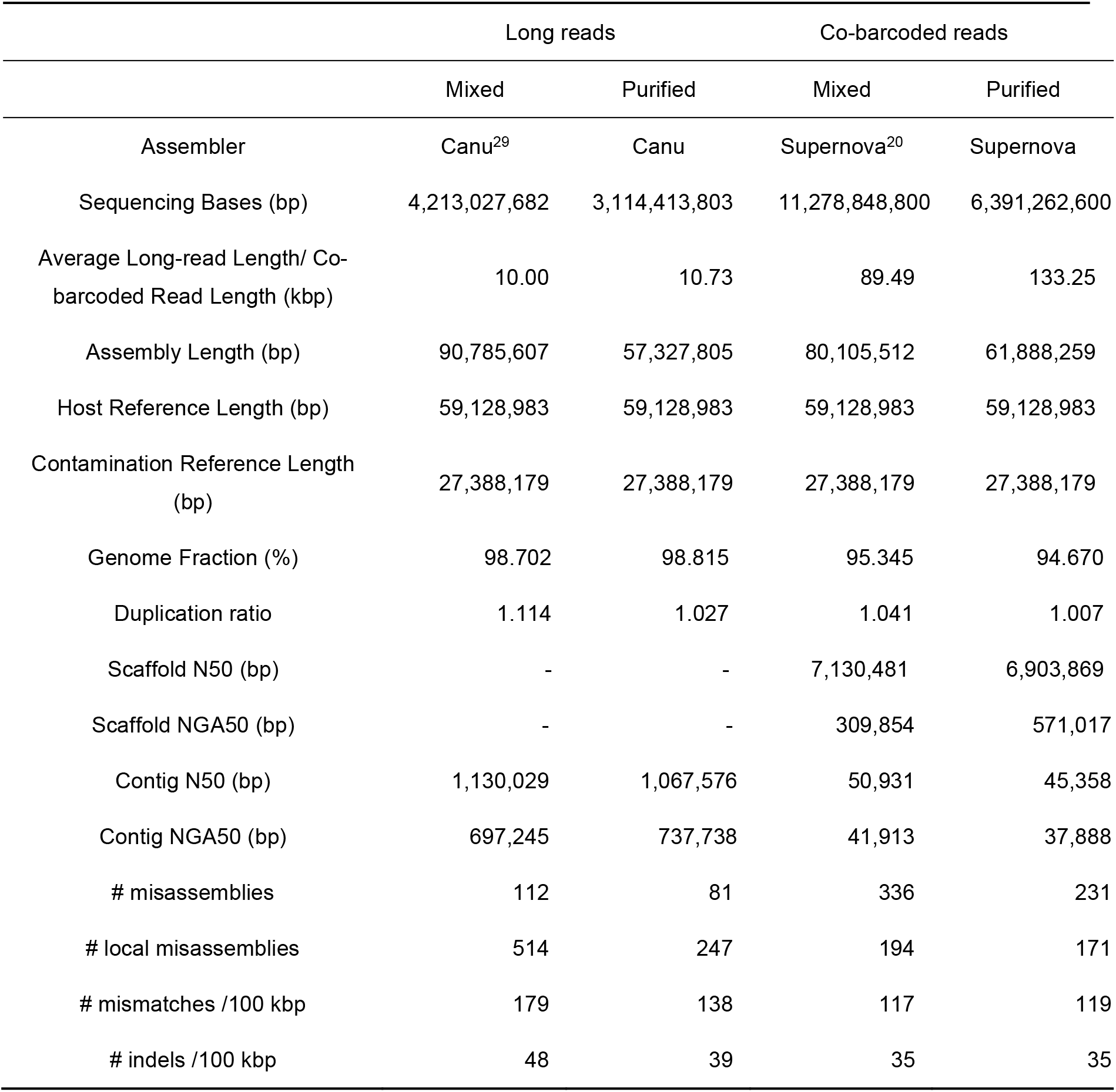
Assembly evaluation for long reads and co-barcoded reads after decontamination.

All long reads or co-barcoded reads were marked by their species names to benchmark the precision and recall rate of clustering. The recall rate is defined as the ratio of the number of clustered host’s reads to the total number of host’s reads in the mixed input. The precision rate is defined as the ratio of the number of correctly assigned reads to the total number of final clustered reads. Note that long fragments with barcodes were clustered according to their features instead of short reads for stLFR data, but the recall and precision rate were calculated based on the co-barcoded reads.

### Generation of parental-unique and shared markers

The contaminants existing with parents were sequenced by the massively parallel sequencing short reads, and mixed with parental reads, respectively. Those combinations of short but accurate reads were used to build the characteristic *k*-mer sets.

We first applied meryl^22^ to build canonical *k*-mer libraries for the two mixed parental-contamination datasets individually. The *k*-mers appearing in paternal group other than maternal were defined as paternal-only makers, while those only existing in maternal group were defined as maternal-only. Meanwhile, those *k*-mers shared by two individuals could be used to filter out sequences from occasional contamination (**Fig. S1**).

### Unsupervised clustering of BGMM

The algorithm of BGMM was selected according to the applicable geometry and running speed from the comparison of the clustering algorithms in scikit-learn^35^. By looking up those characteristic *k*-mers in each single-molecule long read or barcoded long fragment, their *k*-mer densities were obtained and defined as the read feature 1-3. The GC content and canonical 2-mer frequency distribution were defined as the read feature 4-20 (**Table S1**). The preprocessing of PCA decomposed those features into new independent variables in 20-dimensional matrices. The preprocessing of whitening was applied to reduce the informatic redundance. Application Programming Interface from scikit-learn^35^ were used to implement these steps.

### Determination and consensus of best clusters

Based on Algorithm1, the priori host reads were determined by the following formula (*x*l – *threshold*l >0 || *x*2 – *threshold*1 > 0)&& *x*3 – threshold2; > 0 where *x1, x2* refer to parental-only makers, and *x3* refers to shared makers. Then we individually ran the clustering *n* (default 30) times, and selected the best group in each run by the criterion that whether the group contained the most priori high-confident host reads with the smallest variance. The final host group was produced by merging reads in all the best groups and filtering those that occurred less than a frequency threshold (default *n*/5) (**Fig. S2**).

### *k*-mer based evaluation

We used meryl to generate distinct *k*-mers for each species in the mixed sample according to their reference assemblies. The completeness was defined as the ratio of mapped distinct *k*-mer number in the read set or assembly to the total number in the reference. The *k*-mer depth in reads was calculated based on the frequency of mapped *k*-mers.

### Validation of purified assembly

The sequences of the chromosome 19 was extracted from the human reference hs37d5^33^ and used to validate the assemblies. The assembly statistics was reported by QUAST^36^ (version 5.0.2; QUAST, RRID:SCR_001228) with default parameters. Note that contigs in QUAST evaluation was generated by splitting long scaffolds where there is a continuous stretch of ≥ 10 N’s.

## Author contributions

XXX

## Data availability

XXX

## Code availability

All codes and scripts to build *k*-mer sets, extract characteristic features, and classify host reads are available at https://github.com/BGI-Qingdao/Symbiont-Screener.

## Authors’ contributions

XXX

## Acknowledgements

The data that support the findings of this study have been deposited into CNGB Nucleotide Sequence Archive (CNSA)^37^ of China National GeneBank DataBase (CNGBdb)^38^ with accession number CNP0001199.

