## Supplemental Information for "Symbiont-Screener: a reference-free filter to automatically separate host sequences and contaminants for long reads or co-barcoded reads by unsupervised clustering"

**Table S1. The feature composition.**

| Feature name | Feature description |
|---|---|
| *x1* | *k*-mers density of read for *k*-mers from Pat-only *k*-mer library |
| *x2* | *k*-mers density of read for *k*-mers from Mat-only *k*-mer library |
| *x3* | *k*-mers density of read for *k*-mers from Shared *k*-mer library |
| *x4* | GC content of read |
| *x5-x20* | 2-mer frequency of read |

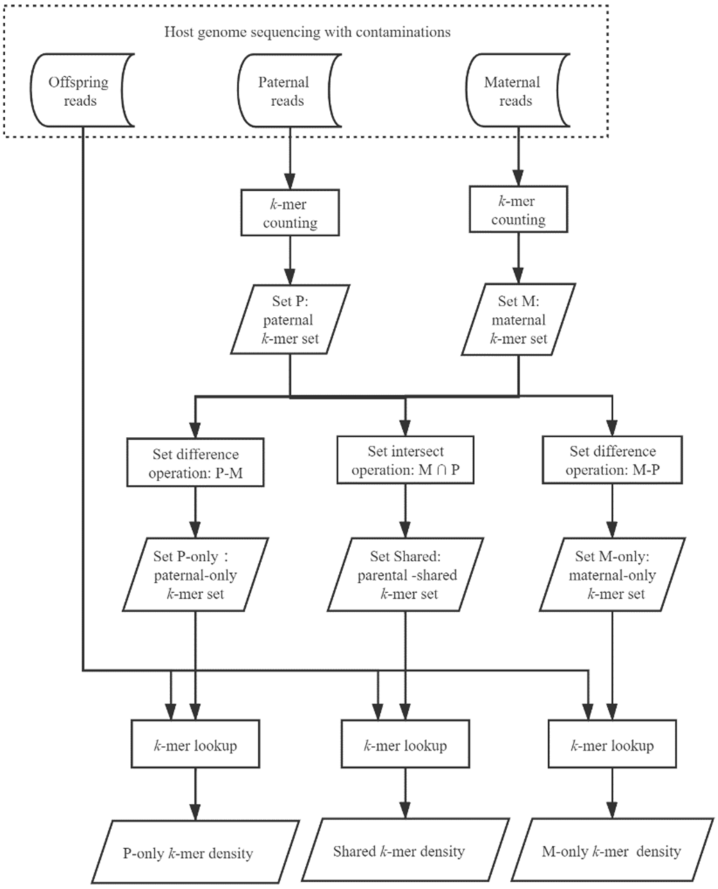


**Fig. S2. The workflow for generating trio *k*-mer density.**

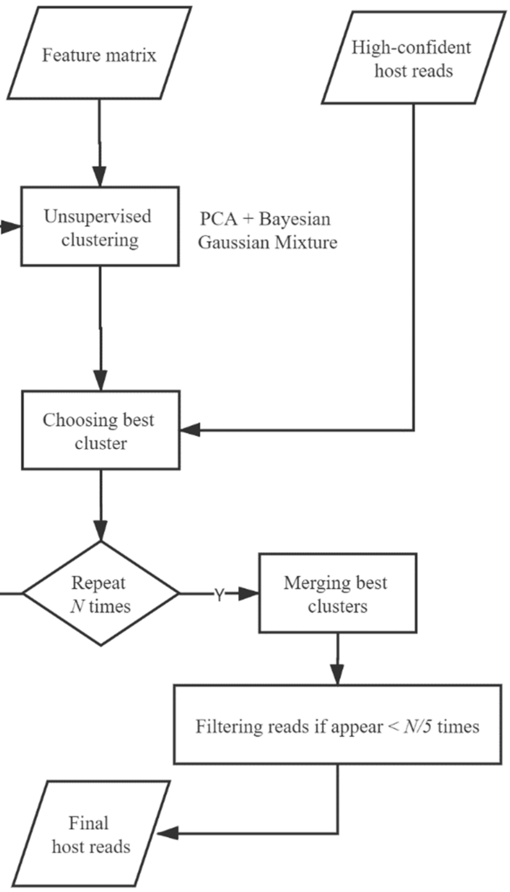


**Fig. S3. The workflow for machine learning part.**
